# Uncertainty alters the balance between incremental learning and episodic memory

**DOI:** 10.1101/2022.07.05.498877

**Authors:** Jonathan Nicholas, Nathaniel D. Daw, Daphna Shohamy

**Affiliations:** Department of Psychology, Columbia University, New York, NY, USA; Mortimer B. Zuckerman Mind, Brain, Behavior Institute, Columbia University, New York, NY, USA; Department of Psychology, Princeton University, Princeton, NJ, USA; Princeton Neuroscience Institute, Princeton University, Princeton, NJ, USA; The Kavli Institute for Brain Science, Columbia University, New York, NY, USA

## Abstract

A key question in decision making is how humans arbitrate between competing learning and memory systems to maximize reward. We address this question by probing the balance between the effects, on choice, of incremental trial-and-error learning versus episodic memories of individual events. Although a rich literature has studied incremental learning in isolation, the role of episodic memory in decision making has only recently drawn focus, and little research disentangles their separate contributions. We hypothesized that the brain arbitrates rationally between these two systems, relying on each in circumstances to which it is most suited, as indicated by uncertainty. We tested this hypothesis by directly contrasting contributions of episodic and incremental influence to decisions, while manipulating the relative uncertainty of incremental learning using a well-established manipulation of reward volatility. Across two large, independent samples of young adults, participants traded these influences off rationally, depending more on episodic information when incremental summaries were more uncertain. These results support the proposal that the brain optimizes the balance between different forms of learning and memory according to their relative uncertainties and elucidate the circumstances under which episodic memory informs decisions.

## Introduction

Effective decision making depends on using memories of past experiences to inform choices in the present. This process has been extensively studied using models of learning from trial-and-error, many of which rely on error-driven learning rules that in effect summarize experiences using a running average^1–3^. This sort of *incremental learning* provides a simple mechanism for evaluating actions without maintaining memory traces of each individual experience along the way, and has rich links to conditioning behavior and putative neural mechanisms for error-driven learning^4^. However, recent findings indicate that decisions may also be guided by the retrieval of individual events, a process often assumed to be supported by *episodic memory*^5–14^. Although theoretical work has suggested a role for episodic memory in initial task acquisition, when experience is sparse^15,16^, the use of episodes may be much more pervasive, as its influence has been detected empirically even in decision tasks that are well-trained and can be solved normatively using incremental learning alone^6,8,10^. The apparent ubiquity of episodic memory as a substrate for decision making raises questions about the circumstances under which it is recruited and the implications for behavior.

How and when episodic memory is used for decisions relates to a more general challenge in cognitive control: understanding how the brain balances competing systems for decision making. An overarching hypothesis is that the brain judiciously adopts different decision strategies in circumstances for which they are most suited; for example, by determining which system is likely to produce the most rewarding choices at the least cost. This general idea has been invoked to explain how the brain Sarbitrates between deliberative versus habitual decisions and previous work has suggested a key role for uncertainty in achieving a balance that maximizes reward^17,18^. Moreover, imbalances in arbitration have been implicated in dysfunction such as compulsion^19,20^, addiction^21,22^, and rumination^23–25^

Here we hypothesized that uncertainty is used for effective arbitration between decision systems and tested this hypothesis by investigating the tradeoff between incremental learning and episodic memory. This is a particularly favorable setting in which to examine this hypothesis due to a rich prior literature theoretically analyzing, and experimentally manipulating, the efficacy of incremental learning in isolation. Studies of this sort typically manipulate the volatility, or frequency of change, of the environment. In line with predictions made by statistical learning models, these experiments demonstrate that when the reward associated with an action is more volatile, people adapt by increasing their incremental learning rates^26–32^. In this case, incrementally constructed estimates reflect a running average over fewer experiences, yielding both less accurate and more uncertain estimates of expected reward. We therefore reasoned that the benefits of incremental learning are most pronounced when incremental estimation can leverage many experiences or, in other words, when volatility is low. By contrast, when the environment is either changing frequently or has recently changed, estimating reward episodically by retrieving a single, well-matched experience should be relatively more favorable.

We tested this hypothesis using a choice task that directly pits these decision systems against one another^11^, while manipulating volatility. In particular, we i) independently measured the contributions of episodic memory vs. incremental learning to choice and ii) altered the uncertainty about incremental estimates using different levels of volatility. Two large online samples of healthy young adults (a primary sample with n=254 and a replication sample with n=223) completed three tasks. The main task of interest combined incremental learning and episodic memory, referred to throughout as the *deck learning and card memory* task (middle panel, **Figure 1A**). On each trial of this task, participants chose between an orange and a blue card and received feedback following their choice. The cards appeared on each trial throughout the task, but their relative value changed over time (**Figure 1B**). In addition to the color of the card, each card also displayed an object. Critically, objects appeared on a card at most twice throughout the task, such that a chosen object could re-appear between 9-30 trials after it was chosen the first time, and would deliver the same reward. Thus, participants could make decisions based on incremental learning of the average value of the orange vs. blue decks, or based on episodic memory for the specific value of an object which they only saw once before. Additionally, participants made choices across two environments: a *high volatility* and a *low volatility* environment. The environments differed in how often reversals in deck value occurred.

**Figure 1.**
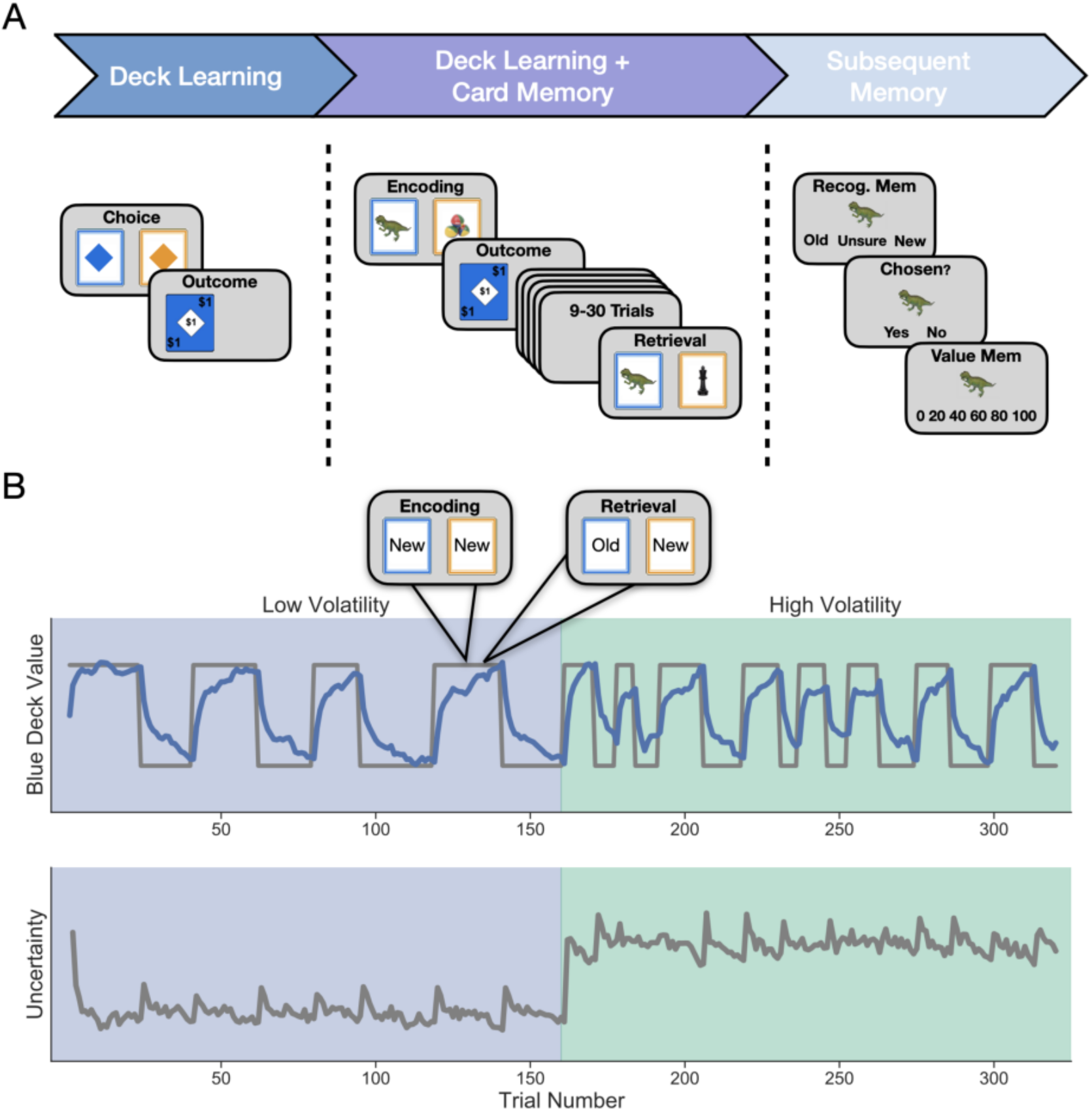
**A) Study Design and Sample Events.** Participants completed three tasks in succession. The first was the *deck learning* task which consisted of choosing between two colored cards and receiving an outcome following each choice. One color was worth more on average at any given timepoint and this mapping changed periodically. Second was the main task of interest, the *deck learning and card memory* task, which followed the same structure as the deck learning task but each card also displayed a trial-unique object. Cards that were chosen could appear a second time in the task after 9-30 trials and, if they re-appeared, were worth the same amount, thereby allowing participants to use episodic memory for individual cards in addition to learning deck value from feedback. Lastly, participants completed a *subsequent memory* task for objects that may have been seen in the deck learning and card memory task. Participants had to indicate whether they recognized an object and, if they did, whether they chose that object. If they responded that they had chosen the object they were then asked if they remembered the value of that object. **B) Uncertainty manipulation within and across environments**. Uncertainty was manipulated by varying the volatility of the relationship between cue and reward over time. Participants completed the task in two environments that differed in their relative volatility. The low volatility environment featured half as many reversals in deck luckiness as the high volatility environment. *Top:* The true value of the blue deck is drawn in gray for an example trial sequence. In blue is estimated blue deck value from the reduced Bayesian model.^30^ Trials featuring objects appeared only in the deck learning and card memory task. *Bottom:* Uncertainty about deck value as estimated by the model is shown in grey. This plot shows relative uncertainty, which is the model’s imprecision in its estimate of deck value.

In addition to the main task, participants also completed two other simple tasks in the experiment. First, participants completed a simple *deck learning* task (left panel, **Figure 1A**) to acclimate them to each environment and quantify the effects of uncertainty. This task included choices between a blue or orange colored diamond on each trial, without any trial-unique objects. Second, after the main task, participants completed a standard *subsequent memory* task (right panel, **Figure 1A**) designed to assess the effects of uncertainty on later episodic memory for objects and value they encountered in the main task.

We predicted that greater uncertainty about incremental values would be related to increased use of episodic memory. The experimental design provided two opportunities to measure the impact of uncertainty both *across* conditions, by comparing between the high and the low volatility environments, and *within* condition, by examining how learning and choices were impacted by each reversal.

## Results

### Episodic memory is used more under conditions of greater uncertainty about deck value

Participants completed two decision making tasks. The *deck learning* task familiarized them with the underlying incremental learning task and established an independent measure of sensitivity to the volatility manipulation. The separate *deck learning and card memory* task measured the additional influence of episodic memory on decisions (**Figure 1**). In the deck learning task participants chose between two decks with expected value that changed periodically across two environments, with one more volatile and the other less volatile. We reasoned that, following each reversal, participants should be more uncertain about deck value and that this uncertainty should reduce over time. Because the more volatile environment featured more reversals, this condition has greater uncertainty overall. In the second deck learning and card memory task, each deck featured cards with trial-unique objects that could re-appear once after being chosen and were worth an identical amount at each appearance. We predicted that decisions would be based more on object value when there was greater uncertainty about deck value. Our logic was that episodic memory should be deployed when incremental learning is inaccurate and unreliable due to frequent or recent change. Thus, we expected choices to be more reliant on episodic memory in the high compared to the low volatility environment and, within an environment, after compared to before reversals.

We first examined whether participants were separately sensitive to each source of value in the deck learning and card memory task: the value of the objects (episodic) and of the decks (incremental). Controlling for average deck value, we found that participants used episodic memory for object value, evidenced by a greater tendency to choose high-valued old objects than low-valued old objects (*β*_*oldvalue*_ = 0.621, 95% *CI* = [0.527, 0.713]; **Figure 2A**). Likewise, controlling for object value, we also found that participants used incrementally learned value for the decks, evidenced by the fact that the higher-valued (lucky) deck was chosen more frequently on trials immediately preceding a reversal (*β*_*t* −4_ = 0.038, 95% *CI* = [−0.038, 0.113]; *β*_*t −*3_ = 0.056, 95% *CI* = [−0.02, 0.134]; *β*_*t −*2_ = 0.088, 95% *CI* = [0.009, 0.166]; *β*_*t* − 1_ = 0.136, 95% *CI* = [0.052, 0.219]; **Figure 2B**), that this tendency was disrupted by the reversals (*β*_*t*+0_ = −0.382, 95% *CI* = [−0.465, −0.296]), and by the quick recovery of performance on the trials following a reversal (*β*_*t+*1_ = −0.175, 95% *CI* = [−0.258, −0.095]; *β*_*t*+2,_ = −0.106, 95% *CI* = [−0.18, −0.029]; *β*_*t*+3_ = −0.084, 95% *CI* = [−0.158, −0.006]; *β*_*t*+4_= 0.129, 95% *CI* = [0.071, 0.184]).

**Figure 2.**
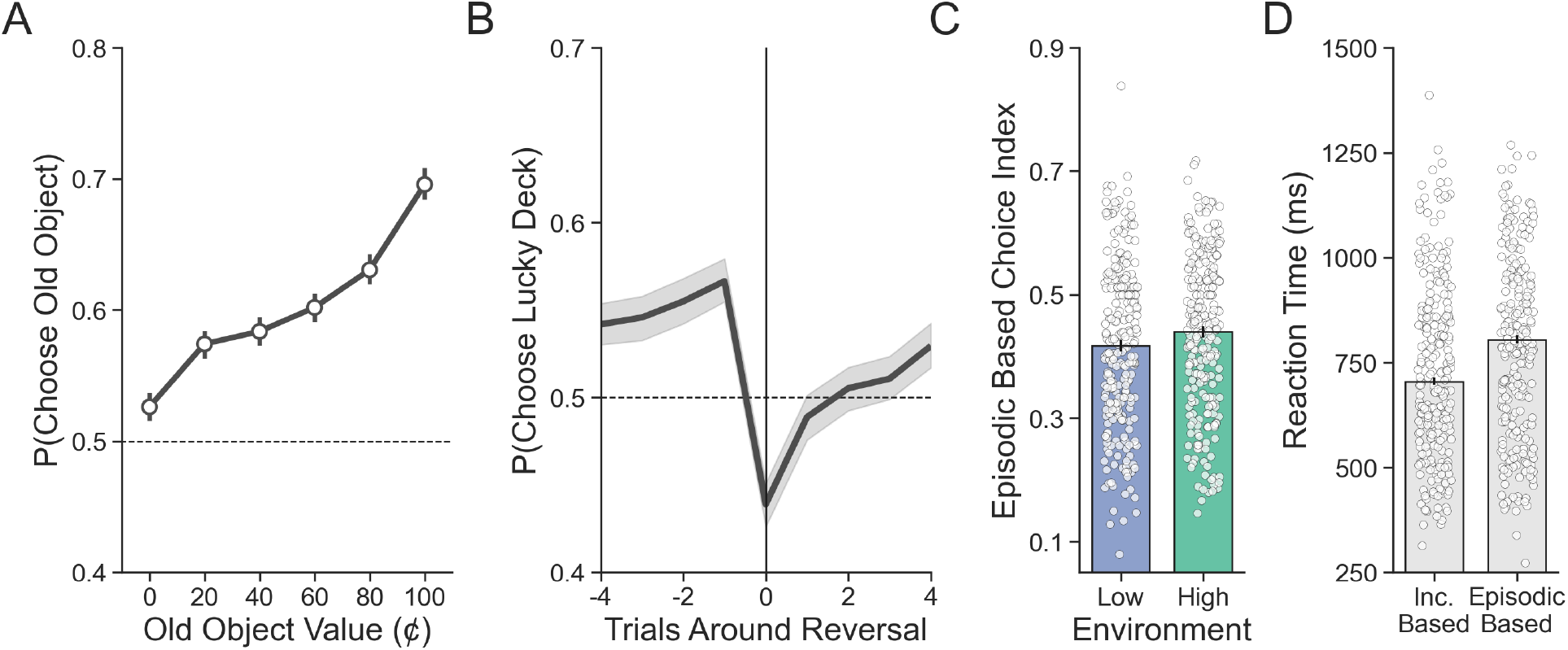
Evaluating the proportion of incremental and episodic choices. **A)** Participants’ choices demonstrate sensitivity to the value of old objects. Group-level averages are shown as points and lines represent 95% confidence intervals. **B)** Reversals in deck luckiness altered choice such that the currently lucky deck was chosen less following a reversal. The line represents the group-level average and the band represents the 95% confidence interval. **C)** On incongruent trials, choices were more likely to be based on episodic memory (e.g. high-valued objects chosen and low-valued objects avoided) in the high compared to the low volatility environment. Averages for individual subjects are shown as points and lines represent the group-level average with a 95% confidence interval. **D)** Median reaction time was longer for incongruent choices based on episodic memory compared to those based on incremental learning.

Having established that both episodic memory and incremental learning guided choices, we next sought to determine the impact of uncertainty on episodic memory for object value by isolating trials on which episodic memory was most likely to be used. To identify reliance on object value, we first focused on trials where the two sources of value information were incongruent: i.e. trials for which the high-value deck featured an old object that was of low value (<50¢) or the low-value deck featured an old object that was of high value (>50¢). We then defined an *episodic based choice index* by considering a choice as episodic if the old object was, in the first case, avoided or, in the second case, chosen. Consistent with our hypothesis, we found greater evidence for episodic choices (as defined this way) in the high volatility environment compared to the low volatility environment (*β*_*Env*_ = 0.094, 95% *CI* = [0.017, 0.17]; **Figure 2C**). Finally, this analysis also gave us the opportunity to test differences in reaction time between incremental and episodic decisions. Decisions based on episodic value took longer (*β*_*EBCI*_ = 38.573, 95% *CI* = [29.703, 47.736]; **Figure 2D**), suggesting that episodic retrieval is more costly in time and perhaps more effortful overall, when compared to relying on cached incremental value.

### Uncertainty in incremental values increases sensitivity to episodic value

To capture uncertainty about deck value on a trial-by-trial basis, we adopted a computational model that tracks uncertainty during learning. We then used this model to test our central hypothesis: that episodic memory is used more when posterior uncertainty about deck value is high.

We began by hierarchically fitting two classes of incremental learning models to the behavior on the deck learning task: a baseline model with a Rescorla-Wagner^2^ style update (RW) and a reduced Bayesian model^30^ (RB) that augments the RW learner with a variable learning rate, which it modulates by tracking ongoing uncertainty about deck value. This approach–which builds on a line of work applying Bayesian learning models to capture trial-by-trial modulation in uncertainty and learning rates in volatile environments^26,27,30,32–34^–allowed us to first assess incremental learning free of any contamination due to competition with episodic memory. We then used the parameters fit to this task for each participant to generate estimates of subjective deck value and uncertainty around deck value, out of sample, in the deck learning and card memory task. These estimates were then used alongside episodic value to predict choices on incongruent trials in the deck learning and card memory task.

We first tested whether participants adjusted their rates of learning in response to uncertainty, both between environments and due to trial-wise fluctuations in uncertainty about deck value. We did this by comparing the ability of each combined choice model to predict participants’ decisions out of sample. To test for effects between environments, we compared models that controlled learning with either a single free parameter (for RW, a learning rate *α*; for RB, a hazard rate *H* capturing the expected frequency of reversals) shared across both environments or models with a separate free parameter for each environment. To test for trial-wise effects within environments, we compared between RB and RW models: while RW updates deck value with a constant learning rate, RB tracks ongoing posterior uncertainty about deck value (called relative uncertainty, RU) and increases its learning rate when this quantity is high.

Participants were both sensitive to the volatility manipulation and incorporated uncertainty into updating their beliefs about deck value. This is indicated by the fact that the RB combined choice model that included a separate hazard rate for each environment (RB2*H*) outperformed both RW models as well as the RB model with a single hazard rate (**Figure 3A**). Further, across the entire sample, participants detected higher levels of volatility in the high volatility environment, as indicated by the generally larger hazard rates recovered from this model in the high compared to the low volatility environment (*H*_*Low*_ = 0.04, 95% *CI* = [0.033, 0.048]; *H*_*High*_= 0.081, 95% *CI* = [0.067, 0.097]; Figure 3B). Next, we examined the model’s ability to estimate uncertainty as a function of reversals in deck luckiness. Compared to an average of the four trials prior to a reversal, RU increased immediately following a reversal and stabilized over time (*β*_*t*+0_ = 0.014, 95% *CI* = [−0.019, 0.048]; *β*_*t*+1_ = −0.242, 95% *CI* = [−0.276, −0.209]; *β*_*t*+2_ = −0.145, 95% *CI* = [−0.178, −0.112]; *β*_*t*+3_ = −0.1, 95% *CI* = [−0.131, −0.07]; *β*_*t*+4_ = −0.079, 95% *CI* = [−0.108, −0.048]; Figure 3C). As expected, RU was also, on average, greater in the high compared to the low volatility environment (*β*_*Env*_ = 0.015, 95% *CI* = [0.012, 0.018]). Lastly, we were interested in assessing the relationship between reaction time and RU, as we expected that higher uncertainty may be reflected in more time needed to resolve decisions. In line with this idea, RU was strongly related to reaction time such that choices made under more uncertain conditions took longer (*β*_*RU*_ = 1.685, 95% *CI* = [0.823, 2.528]).

**Figure 3.**
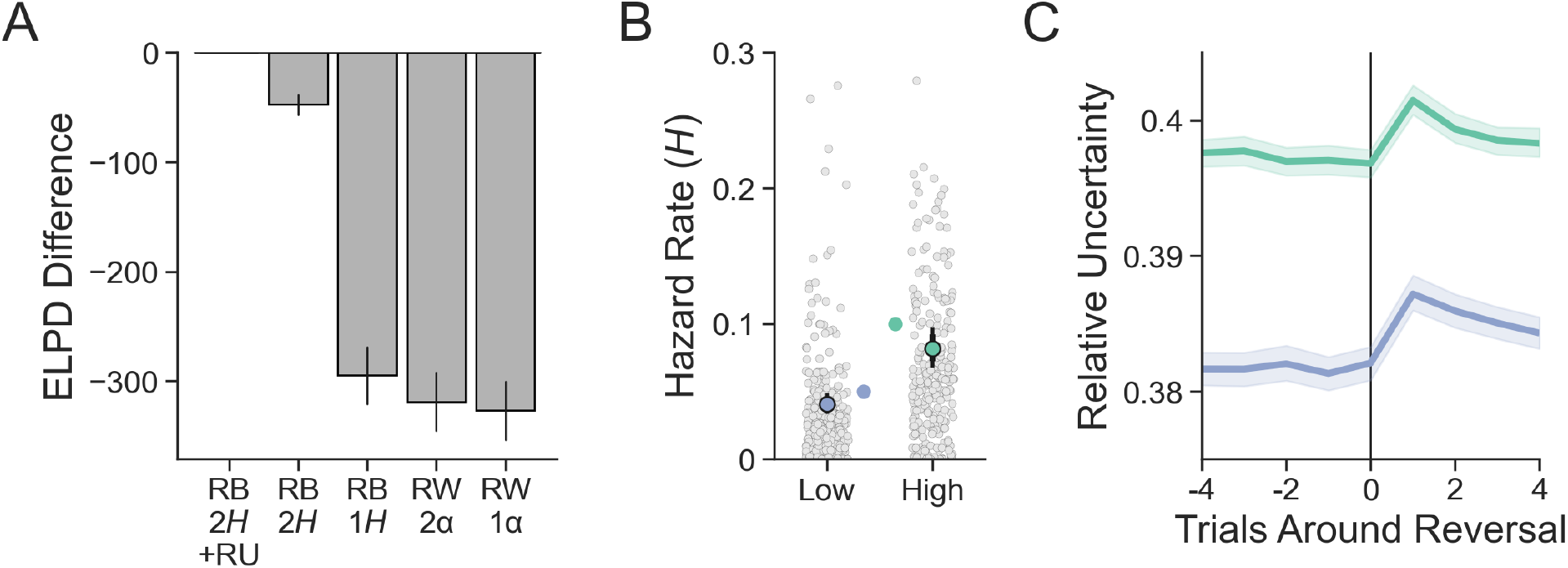
Evaluating model fit and sensitivity to volatility. **A)** Expected log pointwise predictive density from each model was calculated from a 20-Fold leave-N-subjects-out cross validation procedure and is shown here subtracted from the best fitting model. The best fitting model was the reduced Bayesian (RB) model with two hazard rates (2H) and sensitivity to the interaction between old object value and relative uncertainty (RU) in the choice function. Error bars represent standard error around ELPD estimates. **B)** Participants were sensitive to the relative level of volatility in each environment as measured by the hazard rate. Group level parameters are superimposed on individual subject parameters. Wide error bars represent 80% posterior intervals and skinny error bars represent 95% posterior intervals. The true hazard rate for each environment is shown on the interior of the plot. **C)** Relative uncertainty peaks on the trial following a reversal and is greater in the high compared to the low volatility environment. Lines represent group means and bands represent 95% confidence intervals.

Having established that participants were affected by uncertainty around beliefs about deck value, we turned to examine our primary question: whether this uncertainty alters the use of episodic memory in choices. We first examined effects of RU on our episodic choice index, which measures choices consistent with episodic value on trials when it disagrees with incremental learning. This analysis verified that episodic memory was used more on incongruent trial decisions made under conditions of high RU (*β*_*RU*_ = 2.133, 95% *CI* = [0.7, 3.535]; **Figure 4A**). To more directly test the prediction that participants would use episodic memory when uncertainty is high, we included trial-by-trial estimates of RU in the RB2*H* combined choice model, which was augmented with an additional free parameter to capture any change with RU in the effect of episodic value on choice. Formally, this parameter measured an effect of the interaction between these two factors, and the more positive this term the greater the impact of increased uncertainty on the use of episodic memory. This new combined choice model further improved out-of-sample predictions (RB2*H*+RU, **Figure 3A**). As predicted, while both incremental and episodic value were used overall (*β*_*DeckValue*_ = 0.488, 95% *CI* = [0.411, 0.563]; *β*_*Oldvalue*_= 0.141, 95% *CI* = [0.092, 0.19]), episodic value indeed impacted choices more when relative uncertainty was high (*β*_*Oldvalue:RU*_ = 0.091, 95% *CI* = [0.051, 0.13]; **Figure 4B**). This is consistent with our hypothesis that episodic value was relied on more when beliefs about incremental value were uncertain.

**Figure 4.**
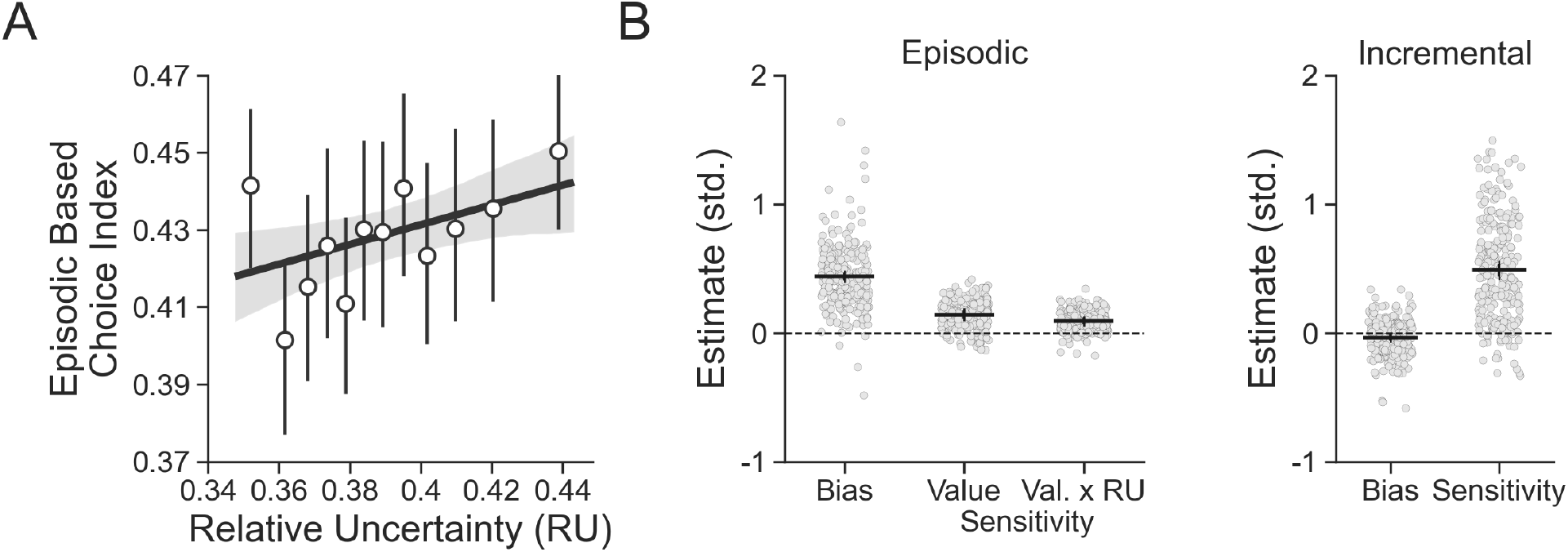
Evaluating effects of sensitivity to uncertainty on episodic choices. **A)** Participants’ degree of episodic-based choice increased with greater RU as predicted by the combined choice model. Points are group means and error bars are 95% confidence intervals. **B)** Estimates from the combined choice model. Participants were biased to choose previously seen objects regardless of their value and were additionally sensitive to their value. As hypothesized, this sensitivity was increased when relative uncertainty was higher. There was no bias to choose one deck color over the other and participants were highly sensitive to estimated deck value. Group level parameters are superimposed on individual subject parameters. Wide error bars represent 80% posterior intervals and skinny error bars represent 95% posterior intervals. Estimates are shown in standard units.

The analyses above focus on uncertainty present at the time of retrieving episodic value because this is what we hypothesized would drive competition in the reliance on either system at choice time. However, in principle, reward uncertainty at the time an object is first encountered might also affect its encoding, and hence its subsequent use in episodic choice when later retrieved^35^. To address this possibility, we looked at the impact of RU resulting from the first time an old object’s value was revealed on whether that object was later retrieved for a decision. Using our episodic based choice index, there was no relationship between the use of episodic memory on incongruent trial decisions and RU at encoding (*β*_*RU*_ = 0.622, 95% *CI* = [−0.832, 2.044]; **Supplementary Figure 5**). Similarly, we also examined effects of trial-by-trial estimates of RU at encoding time in the combined choice model by adding another free parameter that captured change with RU at encoding time in the effect of episodic value on choice. This parameter was added alongside the effect of RU at retrieval time (from the previous analysis). While there was a weak effect on choice (*β*_*Oldvalue:RU*_ = 0.042, 95% *CI* = [0.003, 0.079]; **Supplementary Figure 5**), the inclusion of this parameter did not provide a better fit to subjects’ choices than the combined choice model with only increased sensitivity due to RU at retrieval time (**Supplementary Figure 5**), and this result did not replicate in a separate sample (*β*_*Oldvalue:RU*_ = 0.015, 95% *CI* = [−0.026, 0.057]).

### Episodic and incremental value sensitivity predicts subsequent memory performance

Having determined that decisions depended on episodic memory more when uncertainty about incremental value was higher, we next sought evidence for similar effects on the quality of episodic memory. Episodic memory is, of course, imperfect, and value estimates derived from episodic memory are therefore also uncertain. More uncertain episodic memory should then be disfavored while the influence of incremental value on choice is promoted instead. Although in the present study we did not experimentally manipulate the strength of episodic memory, as our volatility manipulation was designed to affect the uncertainty of incremental estimates, we did measure memory strength in a subsequent memory test. Thus, we predicted that participants who base fewer decisions on object value and more decisions on deck value should have poorer subsequent memory for objects seen in the deck learning and card memory task.

Participants performed well above chance on the test of recognition memory (*β*_0_= 1.887, 95% *CI* = [1.782, 1.989]), indicating a general ability to discriminate objects seen in the main task from those that were new. In line with the idea that episodic memory quality also impacts the relationship between incremental learning and episodic memory, participants with better subsequent recognition memory were more sensitive to episodic value (*β*_*EpSensitivity*_ = 0.373, 95% *CI* = [0.273, 0.478]; Figure 5A), and these same participants were less sensitive to incremental value (*β*_*IncSensitivity*_ = −0.276, 95% *CI* = [−0.383, −0.17]; Figure 5B). This result provides further evidence for a trade-off between episodic memory and incremental learning, and provides preliminary support for a broader version of our hypothesis, which is that uncertainty about value provided by either memory system arbitrates the balance between them.

**Figure 5.**
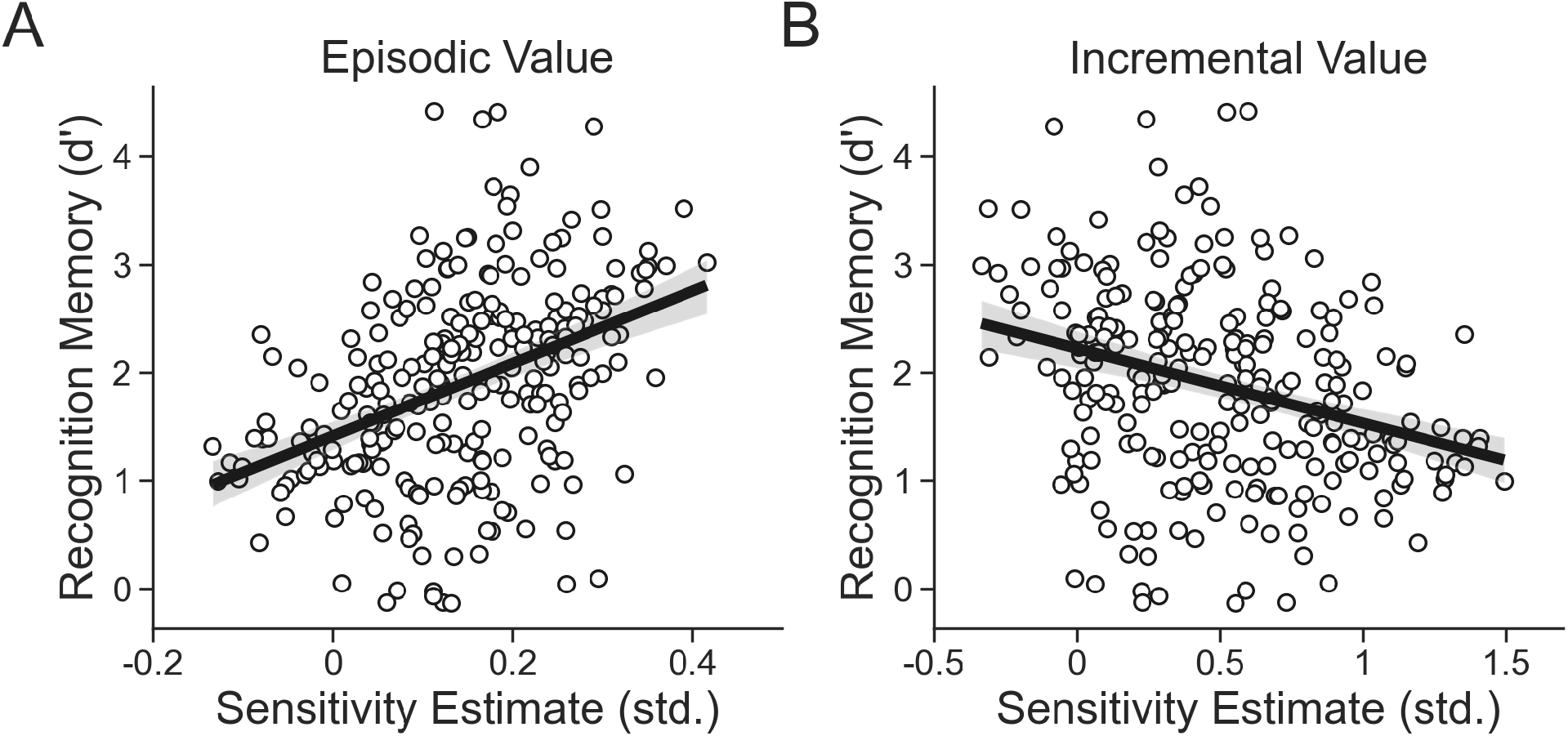
Relationship between choice sensitivity and subsequent memory. **A)** Participants with greater sensitivity to episodic value as measured by random effects in the combined choice model tended to better remember objects seen originally in the card learning and deck memory task. **B)** Participants with greater sensitivity to incremental value tended to have worse memory for objects from the card learning and deck memory task. Points represent individual participants, lines are linear fits and bands are 95% confidence intervals.

### Replication of the main results in a separate sample

We repeated the tasks described above in an independent online sample of healthy young adults (n=223) to test the replicability and robustness of our findings. We replicated all effects of environment and relative uncertainty on episodic-based choice and subsequent memory (see **Supplementary Text** and **Supplementary Figures 1-4** for details).

## Discussion

Research on learning and value-based decision making has focused on how the brain summarizes its experiences by error-driven incremental learning rules that, in effect, maintain the running average of many experiences. While recent work has demonstrated that episodic memory also contributes to value-based decisions^5–14^, many open questions remain about the circumstances under which episodic memory is used. Here we used a task which directly contrasts episodic and incremental influences on decisions and found that participants traded these influences off rationally, relying more on episodic information when incremental summaries were less reliable, i.e. more uncertain and based on fewer experiences. We also found evidence for a complementary modulation of this episodic-incremental balance by episodic memory quality, suggesting that more uncertain episodic-derived estimates may reduce reliance on episodic value. Together, these results indicate that reward uncertainty modulates the use of episodic memory in decisions, suggesting that the brain optimizes the balance between different forms of learning according to volatility in the environment.

Our findings add empirical data to previous theoretical and computational work which has suggested that decision making can greatly benefit from episodic memory for individual estimates when available data are sparse. This most obviously arises early in learning a new task, but also in task transfer, high-dimensional or non-Markovian environments, and (as demonstrated in the current work) during conditions of rapid change^16,36,37^. We investigate these theoretical predictions in the context of human decision making, testing whether humans rely more heavily on episodic memory when incremental summaries comprising multiple experiences are relatively poor. We operationalize this tradeoff in terms of uncertainty, exemplifying a more general statistical scheme for arbitrating between different decision systems by treating them as estimators of action value. There is precedent for this type of uncertainty-based arbitration in the brain, with the most well-known being the tradeoff between model-free learning and model-based learning^17,38^. Control over decision making by model-free and model-based systems has been found to shift in accordance with the accuracy of their respective predictions^18^, and humans adjust their reliance on either system in response to external conditions that provide a relative advantage to one over the other^39–41^. Tracking uncertainty provides useful information about when inaccuracy is expected and helps to maximize utility by deploying whichever system is best at a given time. Our results add to these findings and expand their principles to include episodic memory in this tradeoff.

Indeed, one intriguing possibility is that there is more than just an analogy between the incremental-episodic balance studied here and previous work on model-free versus model-based competition. Incremental error-driven learning coincides closely with model-free learning in other settings^4,17^ and, although it has been proposed that episodic control constitutes a “third way”^16^, it is possible that behavioral signatures of model-based learning might instead arise from episodic control via covert retrieval of individual episodes^15,42–44^, which contain much of the same information as a cognitive map or world model. While the present study assesses single-event episodic retrieval more overtly, it remains an open question for future work the extent to which these same processes, and ultimately the same episodic-incremental tradeoff, might also explain model-based choice as it has been operationalized in other decision tasks. A related line of work has emphasized a similar role for working memory in maintaining representations of individual trials for choice^9,45–47^. Given the capacity constraints of working memory, we think it unlikely that working memory can account for the effects shown here, which involve memory for dozens of trial-unique stimuli maintained over tens of trials.

Further, our findings help to clarify the impacts of uncertainty, novelty, and prediction error on episodic memory more broadly. Recent studies found that new episodes are more likely to be encoded under novel circumstances while prior experiences are more likely to be retrieved when conditions are familiar^11,12,35,48^. Shifts between these states of memory are thought to be modulated by one’s focus on internal or external sources of information^49,50^ and signaled by prediction errors based in episodic memory^51–54^. Relatedly, unsigned prediction errors, which are a marker of surprise, improve later episodic memory^55–58^. Findings have even suggested that states of familiarity and novelty can bias decisions toward the use of single past experiences or not^11,12^. One alternative hypothesis that emerges from this work is that change-induced uncertainty and novelty could exert similar effects on memory, such that novelty signaled by expectancy violations increases encoding in a protracted manner that dwindles as uncertainty is resolved, or the state of the environment becomes familiar. Our results do not support this interpretation. Decisions were guided more by individual memories on more uncertain retrieval trials with little effects of uncertainty at encoding time. It therefore seems likely that uncertainty and novelty operate in concert but remain largely separate concepts, an interpretation supported by recent evidence^59^.

This work raises further questions about the neurobiological basis of memory-based decisions and the role of neuromodulation in signaling uncertainty and aiding memory. In particular, studies have revealed unique functions for norepinephrine (NE) and acetylcholine (ACh) on uncertainty and learning. These findings suggest that volatility, as defined here, is likely to impact the noradrenergic modulatory system, which has been found to signal unexpected changes throughout learning^29,34,60,61^. Noradrenergic terminals densely innervate the hippocampus^62^, and a role for NE in both explicit memory formation^63^ and retrieval^64^ has been posited. Future studies involving a direct investigation of NE or an indirect investigation using pupillometry^29^ may help to isolate its contributions to the interaction between incremental learning and episodic memory in decision making. ACh is also important for learning and memory, as memory formation is facilitated by ACh in the hippocampus, which may contribute to its role in separating and storing new experiences^48,49^. In addition to this role, ACh is heavily involved in incremental learning and has been widely implicated in signaling expected uncertainty, or noise^60,65^. ACh may therefore play an important part in managing the tradeoff between incremental learning and episodic memory. While we held the level of expected uncertainty constant throughout our task, altering this quantity in future work may prove fruitful.

Separately, while in the present study we disadvantaged incremental learning relative to episodic memory, similar predictions about their balance could be made by instead preferentially manipulating episodic memory. There are, for instance, clear theoretical benefits to deploying episodic memory under other task circumstances in which incremental learning is generally ill suited, such as in environments that are high dimensional or require planning far into the future^15^. In principle, individual past experiences can be precisely targeted in these situations depending on the relevance of their features to decisions in the present. Recent advances in computational neuroscience have, for example, demonstrated that artificial agents endowed with episodic memory are able to exploit its rich representation of past experience to make faster, more effective decisions^16,36,37^. While here we provided episodic memory as an alternative source of value to be used in the presence of uncertainty about incremental estimates, future studies making use of paradigms tailored more directly toward episodic memory’s assets will help to further elucidate how and when the human brain recruits episodic memory for decisions.

In conclusion, we have demonstrated that uncertainty induced by volatile environments impacts whether incremental learning or episodic memory is recruited for decisions. Greater uncertainty increased the likelihood that single experiences were retrieved for decision making. This effect suggests that episodic memory aids decision making when simpler sources of value are less accurate. By focusing on uncertainty, our results tie together disparate findings about when episodic memory is recruited for decisions and shed light on the exact circumstances under which the computational expense of episodic memory is worthwhile.

## Materials and Methods

### Experimental Tasks

The primary experimental task used here builds upon a paradigm previously developed by our lab^11^ to successfully measure the relative contribution of incremental and episodic memory to decisions (**Figure 1A**). Participants were told that they would be playing a card game where their goal was to win as much money as possible. Each trial consisted of a choice between two decks of cards that differed based on their color (blue or orange). Participants had two seconds to decide between the decks and, upon making their choice, a green box was displayed around their choice until the full two seconds had passed. The outcome of each decision was then immediately displayed for one second. Following each decision, participants were shown a fixation cross during the intertrial interval period which varied in length (mean = 1.5 seconds, min = 1 seconds, max = 2 seconds). Decks were equally likely to appear on either side of the screen (left or right) on each trial and screen side was not predictive of outcomes. Participants completed a total of 320 trials and were given a 30 second break every 80 trials.

Participants were made aware that there were two ways they could earn bonus money throughout the task, which allowed for the use of incremental and episodic memory respectively. First, at any point in the experiment one of the two decks was “lucky”, meaning that the expected value (*V*) of one deck color was higher than the other (*V*_*lucky*_ =73¢, *V*_*unlucky*_ =27¢). Outcomes ranged from $0 to $1 in increments of 20¢. Critically, the mapping from *V* to deck color underwent an unsignaled reversal periodically throughout the experiment (**Figure 1B**), which incentivized participants to utilize each deck’s recent reward history in order to determine the identity of the currently lucky deck. Each participant completed the task over two environments (with 160 trials in each) that differed in their relative volatility: a low volatility environment with eight *V* reversals, occurring every 20 trials on average, and a high volatility environment with sixteen *V* reversals, occurring every 10 trials on average. Participants were told that they would be playing in two different casinos and that in one casino deck luckiness changed less frequently while in the other deck luckiness changed more frequently. Participants were also made aware of which casino they were currently in by a border on the screen, with a solid black line indicating the low volatility casino and a dashed black line indicating the high volatility casino. Environment order was randomized for each participant.

Second, in order to allow us to assess the use of episodic memory throughout the task, each card within a deck featured an image of a trial-unique object that could re-appear once throughout the experiment after initially being chosen. Participants were told that if they encountered a card a second time it would be worth the same amount as when it was first chosen, regardless of whether its deck color was currently lucky or not. On a given trial *t*, cards chosen once from trials *t* − 9 through *t* − 30 had a 60% chance of reappearing following a sampling procedure designed to prevent each deck’s expected value from becoming skewed by choice, minimize the correlation between the expected value of previously seen cards and deck expected value, and ensure that choosing a previously selected card remained close to 50¢.

Participants also completed a separate decision making task prior to the combined deck learning and card memory task that was identical in design but lacked trial-unique objects on each card. This task, the deck learning task, was designed to isolate the sole contribution of incremental learning to decisions and to allow participants to gain prior experience with each environment’s volatility level. Participants completed the combined deck learning and card memory task immediately following completion of the deck learning task. Instructions were presented immediately prior to each task and participants completed five practice trials and a comprehension quiz prior to starting each.

Following completion of the combined deck learning and card memory task, we tested participants’ memory for the trial-unique objects. Participants completed 80 (up to) three part memory trials. An object was first displayed on the screen and participants were asked whether or not they had previously seen the object and were given five response options: Definitely New, Probably New, Don’t Know, Probably Old, Definitely Old. If the participant indicated that they had not seen the object before or did not know, they moved on to the next trial. If, however, they indicated that they had seen the object before they were then asked if they had chosen the object or not. Lastly, if they responded that they had chosen the object, they were asked what the value of that object was (with options spanning each of the six possible object values between $0-1). Of the 80 trials, 48 were previously seen objects and 32 were new objects that had not been seen before. Of the 48 previously seen objects, half were sampled from each environment (24 each) and, of these, an equal number were taken from each possible object value (with 4 from each value in each environment). As with the decision-making tasks, participants were required to pass a comprehension quiz prior to starting the memory task.

All tasks were programmed using the jsPsych JavaScript library^66^ and hosted on a Google Cloud server running Apache and the Ubuntu operating system. Object images were selected from publicly available stimulus sets^67,68^ for a total of 665 unique objects that could appear in each run of the experiment.

### Participants

A total of 418 participants between the ages of 18 - 35 were recruited for our main sample through Amazon Mechanical Turk using the Cloud Research Approved Participants feature^69^. Recruitment was restricted to the United States and nine dollars of compensation was provided following completion of the 50 minute experiment. Participants were also paid a bonus in proportion to their final combined earnings on both the training task and the combined deck learning and card memory task (total earnings / 100). Before starting each task, all participants were required to score 100% on a quiz that tested their comprehension of the instructions and were made to repeat the instructions until this score was achieved. Informed consent was obtained with approval from the Columbia University Institutional Review Board.

From the initial pool of participants, we excluded those who did not meet our pre-defined performance criteria. Participants were excluded from analysis on the deck learning and card memory task if they i) responded to fewer trials than the group average minus one standard deviation on the deck learning and card memory task, ii) responded faster than the group average minus one standard deviation on this task, or iii) did not demonstrate faster learning in the high compared to the low volatility environment on the independent deck learning task. Our reasoning for this latter decision was that it is only possible to test for effects of volatility on episodic memory recruitment in participants who were sensitive to the difference in volatility between the environments, and it is well-established that a higher learning rate should be used in more volatile conditions^26^. Further, our independent assessment of deck learning was designed to avoid issues of selection bias in this procedure. We measured the effect of environment on learning by fitting a mixed effects logistic regression model to predict if subjects chose the lucky deck up to five trials after a reversal event in the deck learning task. For each subject ***s*** and trial ***t***, this model predicts the probability that the lucky deck was chosen:

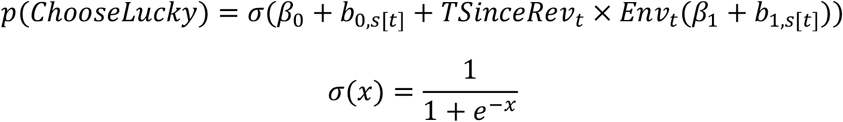

where *β*s are fixed effects, *b*s are random effects, *TSinceRev* is the trial number coded as distance from a reversal event (1-5), and *Env* is the environment a choice was made in coded as −0.5 and 0.5 for the low and high volatility environments respectively. Participants with positive values of *b*_1_ can be said to have chosen the lucky deck more quickly following a reversal in the high compared to the low volatility environment, and we included only these participants in the rest of our analyses. A total of 254 participants survived after applying these criteria.

### Deck Learning and Card Memory Task Behavioral Analysis

We first analyzed the extent to which previously seen (old) objects were used in the combined deck learning and card memory task by fitting the following mixed effects regression model to predict whether an old object was chosen:

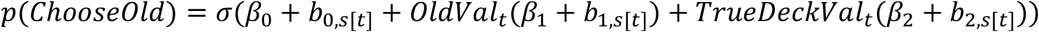

where *OldVal* is the centered value (between −0.5 and 0.5) of an old object. We additionally controlled for the influence of deck value on this analysis by adding a regressor, *TrueDeckVal*, which is the centered true average value of the deck on which each object was shown. Trials not featuring old objects were dropped from this analysis.

We then similarly assessed the extent to which participants engaged in incremental learning overall by looking at the impact of reversals on incremental accuracy directly. To do this, we grouped trials according to their distance from a reversal, up to four trials prior to (*t* = −4: −1), during (*t* = 0), and after (*t* = 1: 4) a reversal occurred. We then dummy coded them to measure their effects on incremental accuracy separately. We also controlled for the influence of old object value in this analysis by including in this regression the coded value of a previously seen object (ranging from 0.5 if the value was $1 on the lucky deck or $0 on the lucky deck to −0.5 if the value was $0 on the lucky deck and $1 on the unlucky deck), for a total of 18 estimated effects:

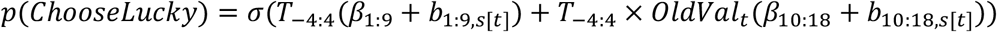

To next focus on whether there was an effect of environment on the extent to which the value of old objects was used for decisions, we restricted all further analyses involving old objects to “incongruent” trials, which were defined as trials on which either the old object was high valued (>50¢) and on the unlucky deck or low valued (<50¢) and on the lucky deck. To better capture participants’ beliefs, deck luckiness was determined by the best-fitting incremental learning model (see next section) rather than using the experimenter-controlled ground truth: whichever deck had the higher model-derived value estimate on a given trial was labeled the lucky deck. Our logic in using only incongruent trials was that choices that stray from choosing whichever deck is more valuable should reflect choices that were based on the episodic value for an object. Lastly, we defined our outcome measure of episodic based choice index (EBCI) to equal 1 on trials where the “correct” episodic response was given (i.e. high valued objects were chosen and low valued object were avoided), and 0 on trials where the “correct” incremental response was given (i.e. the opposite was true). A single mixed effects logistic regression was then used to assess possible effects of environment *Env* on EBCI:

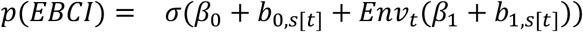

where here *Env* was coded identically to the above analyses.

To assess the effect of episodic-based choices on reaction time (RT), we used the following mixed effects linear regression model:

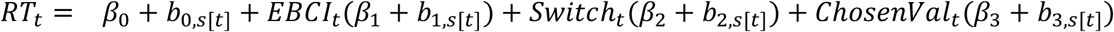

where *EBCI* was coded as −0.5 for incremental-based trials and 0.5 for episodic-based trials. We also included covariates to control for two other possible effects on RT. The first, *Switch*, captured possible RT slowing due to switching from choosing one deck to the other and was coded as −0.5 if a stay occurred and 0.5 if a switch occurred. The second, *ChosenVal*, captured any effects due to the value of the option that may have guided choice, and was set to be the value of the previously seen object on episodic-based trials and the running average true value on incremental-based trials.

For these regression models as well as those described in the following sections, fixed effects are reported in the text as the median of each parameter’s marginal posterior distribution alongside 95% credible intervals, which indicate where 95% of the posterior density falls. Parameter values outside of this range are unlikely given the model, data, and priors. Thus, if the range of likely values does not include zero, we conclude that a meaningful effect was observed.

### Incremental Learning Models

We next assessed the performance of several reinforcement learning models on our task in order to best capture incremental learning. A detailed description of each model can be found in the Supplementary Methods. In brief, these included one model that performed Rescorla-Wagner^2^ style updating with both a single (RW1*α*) and a separate (RW2*α*) fixed learning rate for each environment, and two reduced Bayesian (RB) models^30^ with both a single (RB1*H*) and a separate hazard rate for each environment (RB1*H*). Models were fit to the deck learning task (see **Posterior Inference** and **Supplementary Methods**) and used to generate subject-wise estimates of deck value, and where applicable, uncertainty in the combined deck learning and card memory task.

### Combined Choice Models

After fitting the above hierarchical models to the deck learning task, parameter estimates for each subject were then used to generate trial-by-trial timeseries for deck value and uncertainty (where applicable) throughout performance on the combined deck learning and card memory task. Mixed effects Bayesian logistic regressions for each incremental learning model were then used to capture the effects of multiple memory-based sources of value on incongruent trial choices in this task. For each subject ***s*** and trial ***t***, these models can be written as:

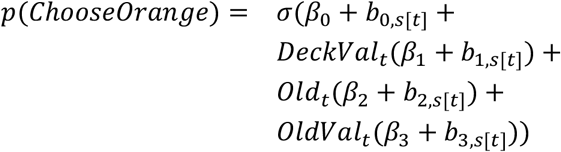

where the intercept captures a bias toward choosing either of the decks regardless of outcome, *DeckVal* is the deck value estimated from each model, the effect of *Old* captures a bias toward choosing a previously seen card regardless of its value, and *OldVal* is the coded value of a previously seen object (ranging from 0.5 if the value was $1 on the orange deck or $0 on the blue deck to −0.5 if the value was $0 on the orange deck and $1 on the blue deck). An additional fifth regression that also incorporated our hypothesized effect of increased sensitivity to old object value when uncertainty about deck value is higher was also fit. This regression was identical to the others but included an additional interaction effect of uncertainty and old object value: *OldVal*_*t*_ ×*Unc*_*t*_*tβ*_4_ + *b*_4,*s*[*t*]_) and used the RB2*H* model’s *DeckVal* estimate alongside its estimate of relative uncertainty (RU) to estimate the effect of *OldVal* ×*Unc*. RU was chosen over CPP because it captures the reducible uncertainty about deck value, which is the quantity we were interested in for this study. Prior to fitting the model, all predictors were z scored in order to report effects in standard units.

### Relative Uncertainty Analyses

We conducted several other analyses that tested effects on or of relative uncertainty (RU) throughout the combined deck learning and card memory task. RU was mean-centered in each of these analyses. First, we assessed separately the effect of RU at retrieval time on EBCI using a mixed effects logistic regression:

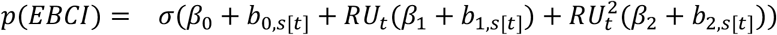

An additional binomial term was included in this model to allow for the possibility that the effect of RU is nonlinear, although this term was found to have no effect. The effect of RU at encoding time was assessed using an identical model but with RU at encoding included instead of RU at retrieval.

Next, to ensure that the RB model captured uncertainty related to changes in deck luckiness, we tested for an effect of environment on RU using a mixed effects linear regression:

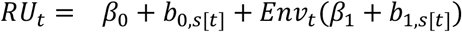

We then also looked at the impact of reversals on RU. To do this, we calculated the difference in RU on reversal trials and up to four trials following a reversal from the average RU on the four trials immediately preceding a reversal. Then, using a dummy coded approach similar to that used for the model testing effects of reversals on incremental accuracy, we fit the following mixed effects linear regression with 5 effects:

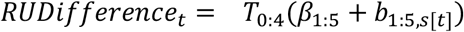

We also assessed the effect of RU on reaction time using another mixed effects linear regression:

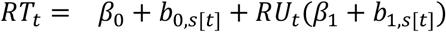

### Subsequent Memory Task Behavioral Analysis

Performance on the subsequent memory task was analyzed in two ways. First, recognition memory was assessed by computing the signal detection metric d prime for each participant adjusted for extreme proportions using a log-linear rule^70^. The relationship with d prime and sensitivity to both episodic value and incremental value was then determined using simple linear regressions of the form *dprime*_*s*_ = *β*_0_ + *Sensitivity*_*s*_(*β*_1_) where *Sensitivity* was either the random effect of episodic value from the combined choice model for each participant or the random effect of incremental value from the combined choice value for each participant.

### Posterior Inference and Model Comparison

Parameters for all incremental learning models were estimated using hierarchical Bayesian inference such that group-level priors were used to regularize subject-level estimates. This approach to fitting reinforcement learning models improves parameter identifiability and predictive accuracy^71^. The joint posterior was approximated using No-U-Turn Sampling^72^ as implemented in stan^73^. Four chains with 2000 samples (1000 discarded as burn-in) were run for a total of 4000 posterior samples per model. Chain convergence was determined by ensuring that the Gelman-Rubin statistic 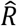 was close to 1. A full description of the parameterization and choice of priors for each model can be found in the **Supplementary Methods**. All regression models were fit using No-U-Turn Sampling in Stan with the same number of chains and samples. Default weakly-informative priors implemented in the rstanarm package^74^ were used for each regression model. Model fit for the combined choice models was assessed by separating each dataset into 20 folds and performing a cross validation procedure by leaving out N/20 subjects per fold where N is the number of subjects in each sample. The expected log pointwise predictive density (ELPD) was then computed and used as a measure of out-of-sample predictive fit for each model.

### Replication

We identically repeated all procedures and analyses applied to the main sample on an independently collected replication sample. A total of 401 participants were again recruited through Amazon Mechanical Turk and 223 survived exclusion procedures carried out identically to those used for the main sample.

### Citation race and gender diversity statement

The gender balance of papers cited within this work was quantified using databases that store the probability of a first name being carried by a woman. Excluding self-citations to the first and last authors of the current paper, the gender breakdown of our references is 12.16% woman(first)/woman(last), 6.76% man/woman, 23.44% woman/man, and 57.64% man/man. This method is limited in that a) names, pronouns, and social media profiles used to construct the databases may not, in every case, be indicative of gender identity and b) it cannot account for intersex, non-binary, or transgender people. Second, we obtained predicted racial/ethnic category of the first and last author of each reference using databases that store the probability of a first and last name being carried by an author of color. By this measure (and excluding self-citations), our references contain 9.55% author of color (first)/author of color(last), 19.97% white author/author of color, 22.7% author of color/white author, and 47.78% white author/white author. This method is limited in that a) using names and Florida Voter Data to make the predictions may not be indicative of racial/ethnic identity, and b) it cannot account for Indigenous and mixed-race authors, or those who may face differential biases due to the ambiguous racialization or ethnicization of their names.

## Data Availability

All code, data, and software needed to reproduce the manuscript can be found here: https://codeocean.com/capsule/2024716/tree/v1

## Contributions

J.N., D.S., and N.D.D. designed the study. J.N. conducted the experiments and analyzed the data. J.N., D.S., and N.D.D. wrote the manuscript.

## Acknowledgements

The authors thank Sam Gershman, Raphael Gerraty, Camilla van Geen, Mariam Aly and members of the Shohamy Lab for insightful discussion and conversations. Support was provided by the NSF Graduate Research Fellowship (J.N.; award # 1644869), the NSF (D.S., N.D.; award # 1822619), the NIMH/NIH (D.S., N.D.; award # MH121093), and the Templeton Foundation (D.S. grant #60844).

## Supplementary Text

**Supplementary Figure 1.**
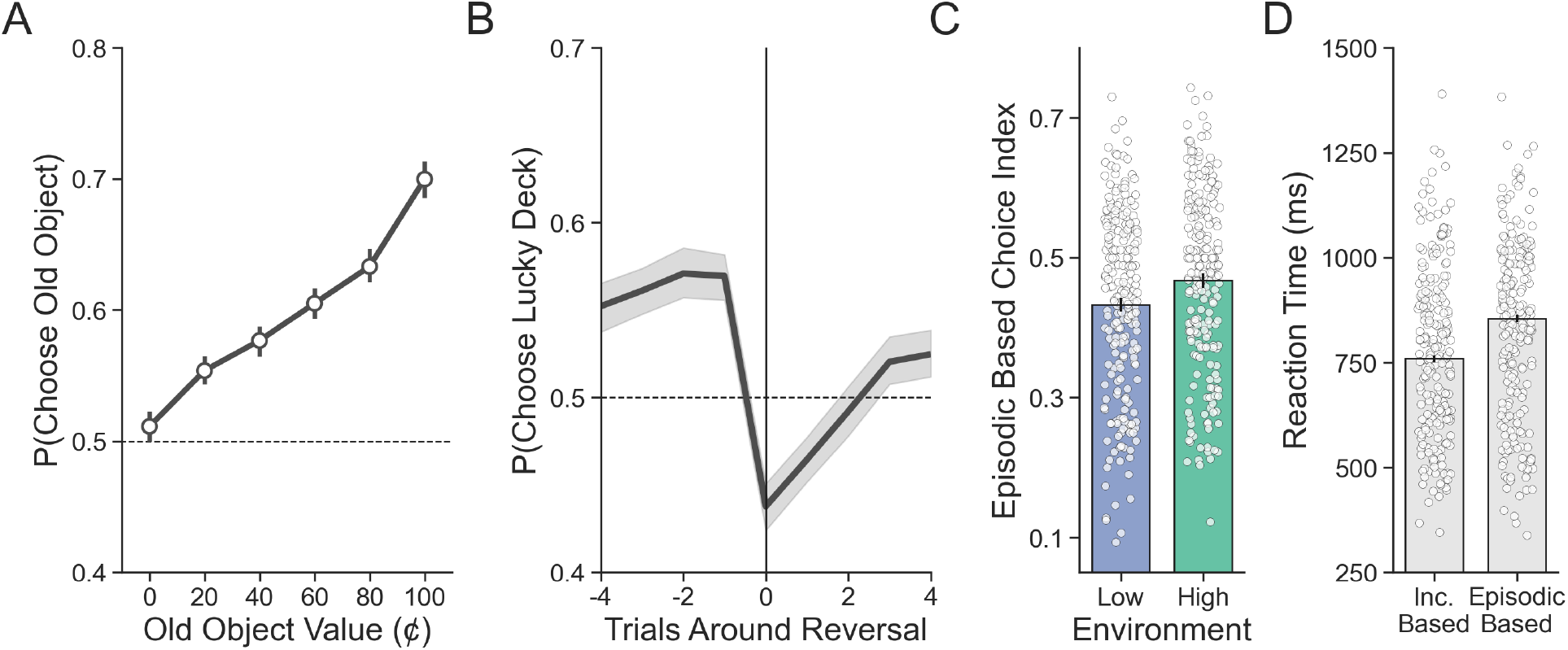
Recreation of Figure 2 in the main text using the replication dataset. **A)** Participants’ choices demonstrate sensitivity to the value of old objects. **B)** Reversals in deck luckiness altered choice such that the currently lucky deck was chosen less following a reversal. **C)** On incongruent trials, choices were more likely to be based on episodic memory in the high compared to the low volatility environment. **D)** Reaction time was longer for incongruent choices based on episodic memory compared to those based on incremental learning.

**Supplementary Figure 2.**
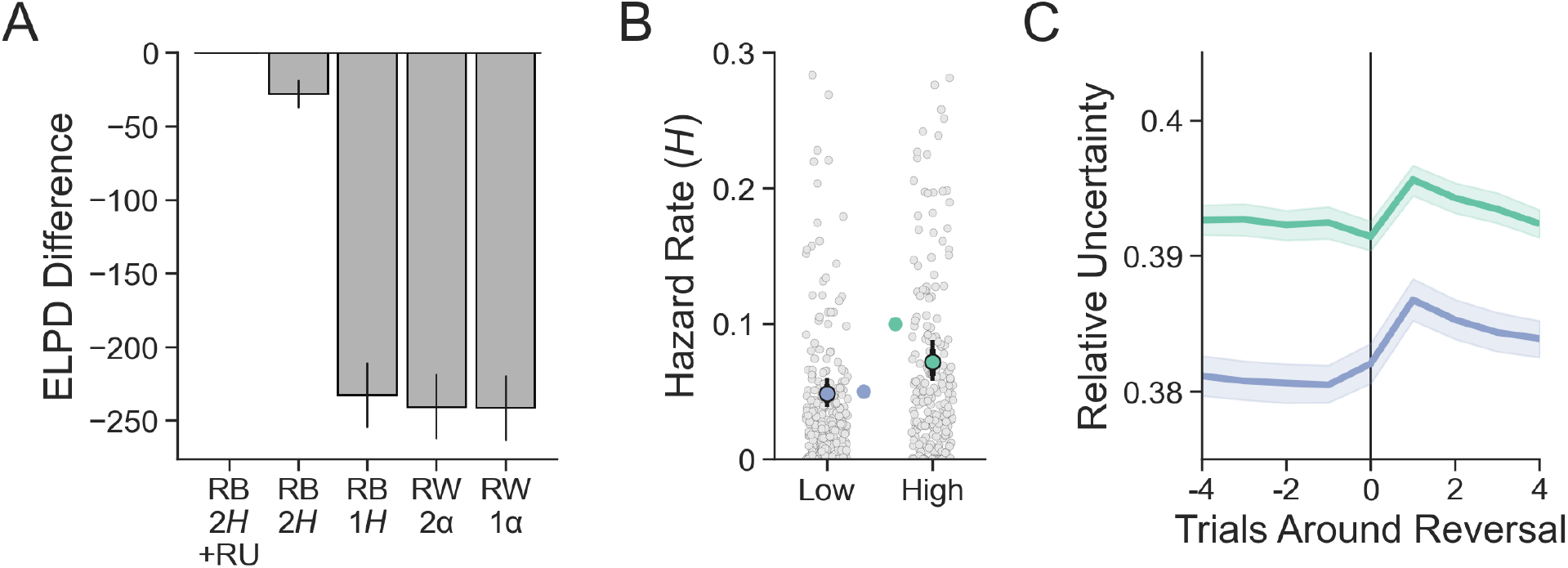
Recreation of Figure 3 in the main text using the replication dataset. **A)** The best fitting model was again the reduced Bayesian (RB) model with two hazard rates (2H) and sensitivity to the interaction between old object value and relative uncertainty (RU) in the choice function. **B)** Participants were affected by the relative level of volatility in each environment as measured by the hazard rate. Group level parameters are superimposed on individual subject parameters. **C)** Relative uncertainty peaks on the trial following a reversal and is greater in the high compared to the low volatility environment.

**Supplementary Figure 3.**
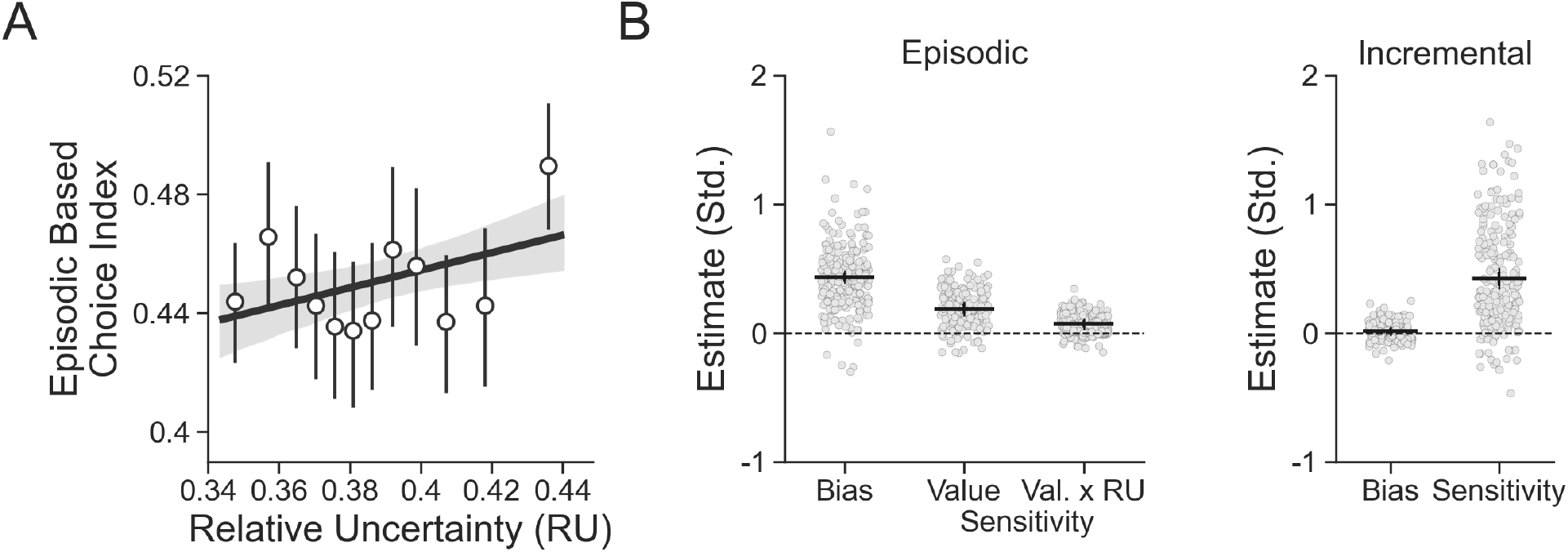
Recreation of Figure 4 in the main text using the replication dataset. **A)** Participants’ degree of episodic-based choice increases with greater RU. **B)** Estimates from the combined choice model. Participants were biased to choose previously seen objects regardless of their value and were additionally sensitive to their value. As hypothesized, this sensitivity was increased when relative uncertainty was higher. There was no bias to choose one deck color over the other and participants were highly sensitive to estimated deck value.

**Supplementary Figure 4.**
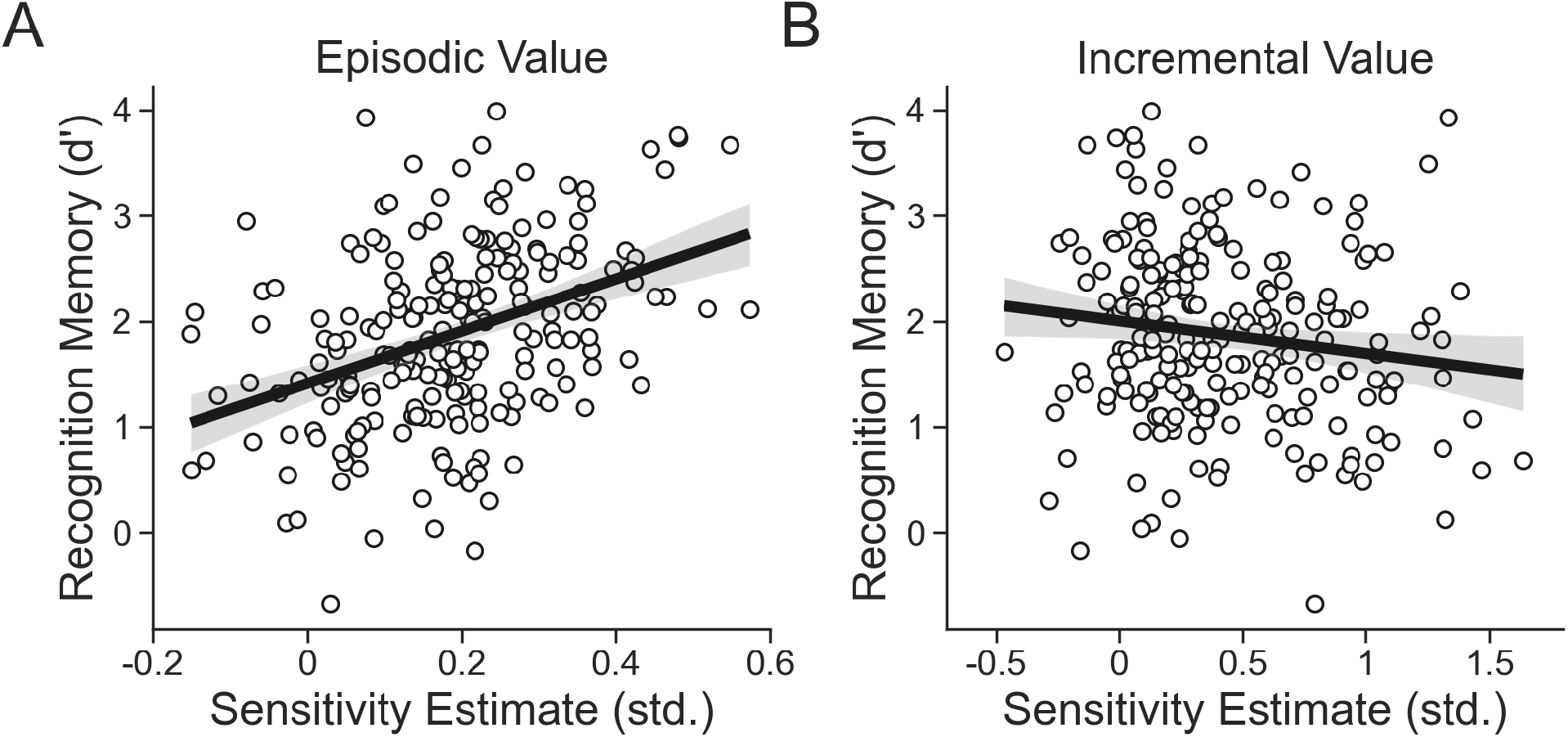
Recreation of Figure 5 in the main text using the replication dataset. **A)** Participants with greater sensitivity to episodic value tended to better remember objects from the deck learning and card memory task. **B)** Participants with greater sensitivity to incremental value tended to have worse memory for objects from the card learning and deck memory task.

**Supplementary Figure 5.**
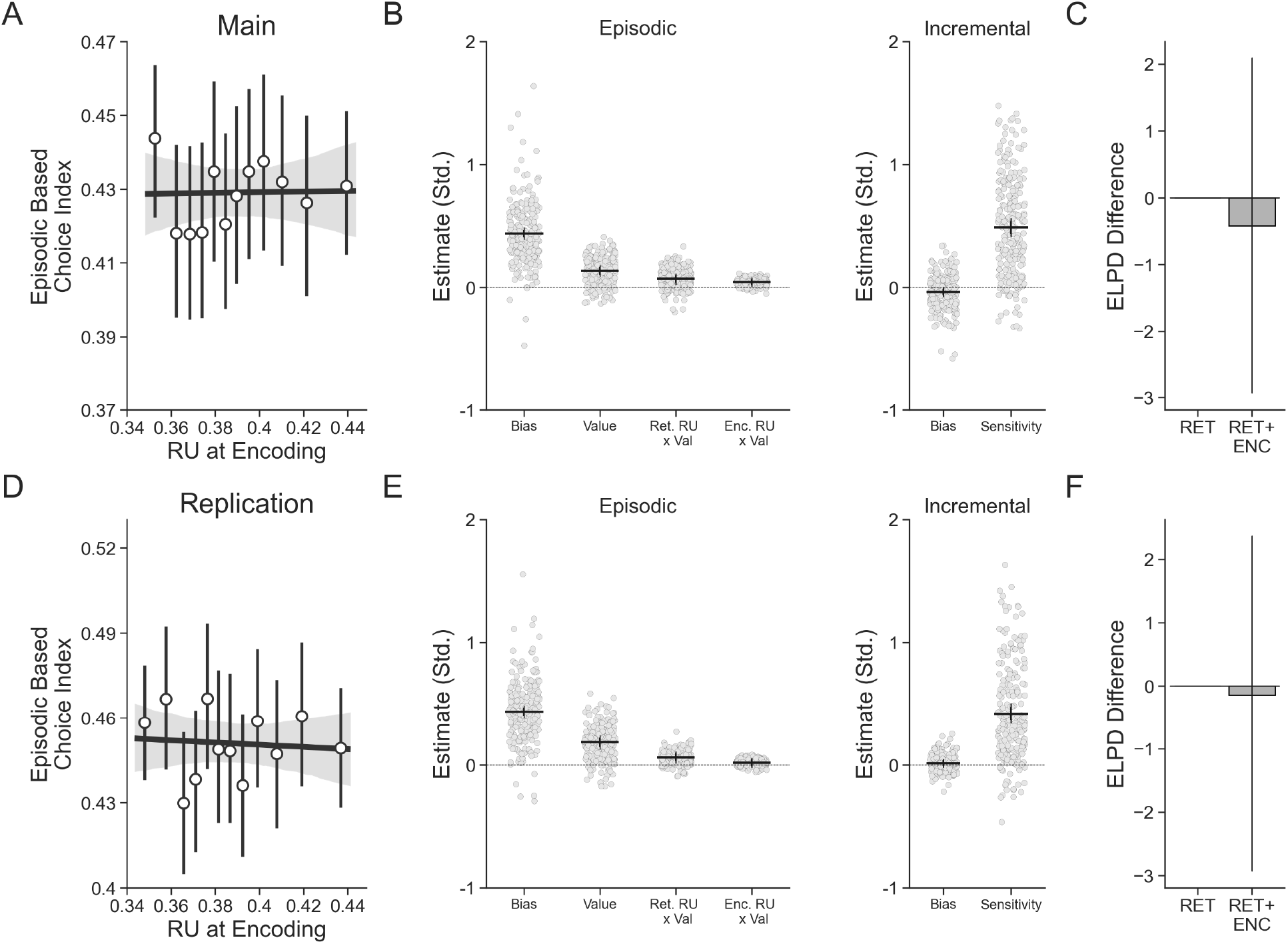
Results of relative uncertainty (RU) at encoding time on episodic-based choice in the main (**A**,**B**,**C**) and replication (**D**,**E**,**F**) sample. **A)** There was no relationship between RU at encoding and the degree to which participants based decisions on episodic value. **B)** Estimates from the combined choice model including both effects of RU at retrieval time and RU at encoding time. Relative to the effect of the interaction between RU at retrieval time and old object value, the equivalent effect for RU at encoding time was small in the main sample. **C)** Expected log pointwise predictive density for the combined choice model including only an effect of the interaction between RU at retrieval time and old object value (presented in the text) and the model also including the interaction between RU at encoding time and old object value. Including RU at encoding time did not improve model performance. **D)** There was again no relationship between RU at encoding and episodic-based choice in the replication sample. **E)** In the replication sample, there was no effect of the interaction between RU at encoding and old object value on choice behavior. **F)** Including RU at encoding time again did not improve model performance in the replication sample.

### Replication Results

Here we repeat and describe all analyses reported in the main text with replication sample. All results are reported in the same order as in the main text.

### Episodic memory is used more under conditions of greater uncertainty

Participants in the replication sample were substantially more likely to chose high-valued old objects compared to low-valued old objects (*β*_*OldValue*_ = 0.723, 95% *CI* = [0.624, 0.827]; **Supplementary Figure 1A**). Participants also altered their behavior in response to reversals in deck value. The higher-valued (lucky) deck was chosen more frequently on trials immediately preceding a reversal (*β*_*t*−4_ = 0.095, 95% *CI* = [0.016, 0.176]; *β*_*t*−3_ = 0.128, 95% *CI* = [0.047, 0.213]; *β*_*t*−2_ = 0.168, 95% *CI* = [0.085, 0.251]; *β*_*t* − 1_ = 0.161, 95% *CI* = [0.075, 0.25]; **Supplementary Figure 1B**). This tendency was then disrupted by trials on which a reversal occurred (*β*_*t*=0_ = −0.373, 95% *CI* = [−0.464, −0.286]), with performance quickly recovering as the newly lucky deck became chosen more frequently on the trials following a reversal (*β*_*t*+1_ =−0.256, 95% *CI* = [−0.337, −0.175]; *β*_*t*+2_ = −0.144, 95% *CI* = [−0.22, −0.064]; *t* + 3: *β*_*t*+3_ = −0.024, 95% *CI* = [−0.102, 0.053]; *β*_*t*+4_ = 0.113, 95% *CI* = [0.055, 0.174]). Thus, participants in the replication sample were also sensitive to reversals in deck value, thereby indicating that they engaged in incremental learning throughout the task.

Participants in the replication sample also based more decisions on episodic value in the high volatility environment compared to the low volatility environment (*β*_*Env*_ = 0.145, 95% *CI* = [0.063, 0.229]; **Supplementary Figure 1C**). Furthermore, decisions based on episodic value again took longer (*β*_*EBCI*_ = 41.38, 95% *CI* = [30.823, 51.707]; **Supplementary Figure 1D**).

### Uncertainty increases sensitivity to episodic value

In the replication sample, the reduced Bayesian model with two hazard rates was again the best fitting model (**Supplementary Figure 2A**). Participants detected higher levels of volatility in the high compared to the low volatility environment, as indicated by the generally larger hazard rates recovered from the high compared to the low volatility environment (*β*_*Low*_ = 0.048, 95% *CI* = [0.038, 0.06]; *β*_*High*_ = 0.071, 95% *CI* = [0.058, 0.088]; **Supplementary Figure 2B**). Compared to an average of the four trials prior to a reversal, RU also increased immediately following a reversal and stabilized over time (*β*_*t*=0_ = 0.021, 95% *CI* = [−0.014, 0.056]; *β*_*t*+1_ = −0.22, 95% *CI* = [−0.253, −0.185]; *β*_*t*+2_ = −0.144, 95% *CI* = [−0.178, −0.11]; *β*_*t*+3_ = −0.098, 95% *CI* = [−0.129, −0.064]; *β*_*t*+4_ = −0.05, 95% *CI* = [−0.083, −0.019]; **Supplementary Figure 2C**). RU was again also, on average, greater in the high compared to the low volatility environment (*β*_*Env*_ = 0.01, 95% *CI* = [0.007, 0.013]) and related to reaction time such that choices made under more uncertain conditions took longer (*β*_*RU*_ = 1.364, 95% *CI* = [0.407, 2.338]).

Episodic memory was also used more on incongruent trial decisions made under conditions of high RU (*β*_*RU*_ = 2.718, 95% *CI* = [1.096, 4.436]; **Supplementary Figure 3A**). We again fit the combined choice model to the replication sample and found the following. Participants again used both sources of value throughout the task: both deck value as estimated by the model (*β*_*DeckValue*_ = 0.42, 95% *CI* = [0.336, 0.505]; **Supplementary Figure 3B**) and the episodic value from old objects (*β*_*OldValue*_ = 0.188, 95% *CI* = [0.13, 0.245]) strongly impacted choice. Lastly, episodic value again impacted choices more when relative uncertainty was high (*β*_*OldValue:RU*_ = 0.069, 95% *CI* = [0.024, 0.113]).

Finally, there was again no relationship between the use of episodic memory on incongruent trial decision and RU at encoding (*β*_*RU*_ = 0.99, 95% *CI* = [−0.642, 2.576]; **Supplementary Figure 5**).

Unlike in the main sample, however, including a sixth parameter to assess increased sensitivity to old object value due to RU at encoding time did not have an effect in the combined choice model (*β*_*OldValue:RU*_ = 0.015, 95% *CI* = [−0.026, 0.057]; **Supplementary Figure 5**), which is also reported in the main text. As with the main sample, including this parameter did not provide a better fit to subjects’ choices than the combined choice model with only increased sensitivity due to RU at retrieval time.

### Episodic and Incremental value sensitivity predicts subsequent memory performance

Participants in the replication sample again performed well above chance on the test of recognition memory (*β*_0_ = 1.874, 95% *CI* = [1.772, 1.977]). Participants with better subsequent recognition memory were again more sensitive to episodic value (*β*_*EpSensitivity*_ = 0.334, 95% *CI* = [0.229, 0.44]; **Supplementary Figure 4A**), and these same participants were again less sensitive to incremental value (*β*_*IncSensitivity*_ = −0.124, 95% *CI* = [−0.238, −0.009]; **Supplementary Figure 4B**).

## Supplementary Methods

### Description of Incremental Learning Models

#### Rescorla Wagner (RW)

The first model we considered was a standard model-free reinforcement learner that assumes a stored value (*Q*) for each deck is updated over time. *Q* is then referenced on each decision in order to guide choices. After each outcome *o*_*t*_, the value for the orange deck *Q*_*0*_ is updated according to the following rule^1^ if the orange deck chosen:

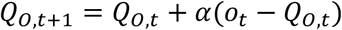

And is not updated if the blue deck is chosen:

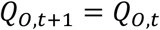

Likewise, the value for the blue deck *Q*_*B*_ is updated equivalently. Large differences between estimated value and outcomes therefore have a larger impact on updates, but the overall degree of updating is controlled by the learning rate, *α*. Two versions of this model were fit, one with a single learning rate (RW1*α*), and one with two learning rates (RW2*α*), *α*_*low*_ or *α*_*high*_ depending on which environment the current trial was completed in. These parameters are constrained to lie between 0 and 1. A separate learning rate was used for each environment in the (RW2*α*) version to capture the well-established idea that a higher learning rate should be used in more volatile conditions^2^.

#### Reduced Bayesian (RB)

The second model we considered was the reduced Bayesian (RB) model developed by Nassar and colleagues^3^. This model tracks and updates its belief that the orange deck is lucky based on trialwise outcomes, *o*_*t*_, using the following prediction error-based update:

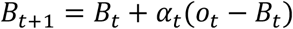

This update is identical to that used in the RW model, however the learning rate *α*_*t*_ is itself updated following each outcome according to the following rule:

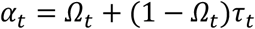

where *Ω*_*t*_ is the probability that a change in deck luckiness has occurred on the most recent trial (the change point probability or CPP) and *τ*_*t*_ is the imprecision in the model’s belief about deck value (the relative uncertainty or RU). The learning rate therefore increases whenever CPP or RU increase. CPP can be written as:

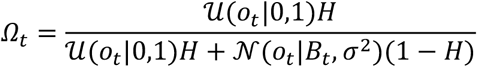

where *H* is the hazard rate or probability of a change in deck luckiness. Two versions of this model were fit, one with a single hazard rate (RB1*H*), and one with two hazard rates (RB2*H*), *H*_*Low*_ and *H*_*High*_, depending on the environment the current trial was completed in. In this equation, the numerator represents the probability that an outcome was sampled from a new average deck value, whereas the denominator indicates the combined probability of a change and the probability that the outcome was generated by a Gaussian distribution centered around the most recent belief about deck luckiness and the variance of this distribution, *σ*^2^. Because CPP is a probability, it is constrained to lie between 0 and 1. In our implementation, *H* was a free parameter (see Posterior Inference section below) and *Ω*_1_ was initialized to 1.

RU, which is the uncertainty about deck value relative to the amount of noise in the environment, is quite similar to the Kalman gain used in Kalman filtering^4^:

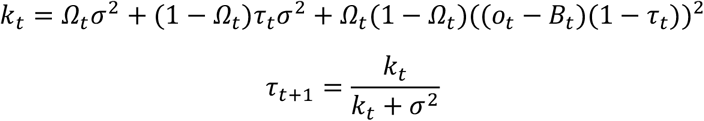

where *σ*^2^ is the observation noise and was here fixed to the true observation noise (0.33). *k*_*t*_ consists of three terms: the first is the variance of the deck value distribution conditional on a change point, the second is the variance of the deck value distribution conditional on no change, and the third is the variance due to the difference in means between these two distributions. These terms are then used in the equation for *τ*_*t*+ 1_ to provide the uncertainty about whether an outcome was due to a change in deck value or the noise in observations that is expected when a change point has not occurred. Because this model does not follow the two-armed bandit assumption of our task (that is, that outcomes come from two separate decks), all outcomes were coded in terms of the orange deck. For example, this means that an outcome worth $1 on the orange deck is treated the same as an outcome worth $0 on the blue deck by this model. While this description represents a brief overview of the critical equations of the reduced Bayesian model, a full explanation can be found in Nassar et al., 2010^3^.

#### Softmax Choice

All incremental learning models were paired with a softmax choice function in order to predict participants’ decisions on each trial:

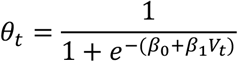

where θ_*t*_ is the probability that the orange deck was chosen on trial *t*. This function also consists of two inverse temperature parameters: *β*_0_ to model an intercept and *β*_1_ to model the slope of the decision function related to deck value. The primary difference for each model was how *V*_*t*_ is computed: RW (*V*_*t*_ = *Q*_0,*t*_ – *Q*_*B,t*_); RB (*V*_*t*_ = *B*_*t*_). In each of these cases, a positive *V*_*t*_ indicates evidence that the orange deck is more valuable while a negative *V*_*t*_ indicates evidence that the blue deck is more valuable.

#### Posterior Inference

For all incremental learning models, the likelihood function can be written as:

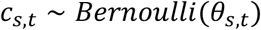

where *c*_*s,t*_ is 1 if subject *s* chose the orange deck on trial *t* and 0 if blue was chosen. Following the recommendations of Gelman and Hill, 2006^5^ and van Geen and Gerraty, 2021^6^, *β*_*s*_ is drawn from a multivariate normal distribution with mean vector *µ*_*β*_ and covariance matrix *Σ*_*β*_:

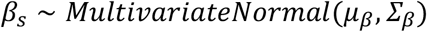

where *Σ*_*β*_ is decomposed into a vector of coefficient scales *τ*_*β*_ and a correlation matrix *Ω*_*β*_ via:

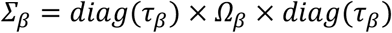

Weakly-informative hyperpriors were then set on the hyperparameters *µ*_*β*_, *Ω*_*β*_ and *τ*_*β*_:

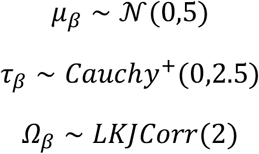

These hyperpriors were chosen for their respective desirable properties: the half cauchy is bounded at zero and has a relatively heavy tail which is useful for scale parameters, the LKJ prior with shape = 2 concentrates some mass around the unit matrix thereby favoring less correlation^7^, and the normal is a standard choice for regression coefficients.

Because sampling from heavy tailed distributions like the Cauchy is difficult for Hamiltonian Monte Carlo^8^, a reparameterization of the Cauchy distribution was used here. *τ*_*β*_ was thereby defined as the transform of a uniformly distributed variable *τ*_*β* __ *u* using the Cauchy inverse cumulative distribution function such that:

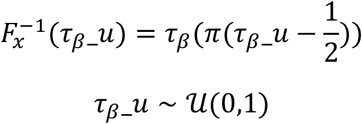

In addition, a multivariate non-centered parameterization specifying the model in terms of the Cholesky factorized correlation matrix was used in order to shift the data’s correlation with the parameters to the hyperparameters, which increases the efficiency of sampling the parameters of hierarchical models^8^. The full correlation matrix Ω_*β*_ was replaced with a Cholesky factorized parameter 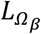 such that:

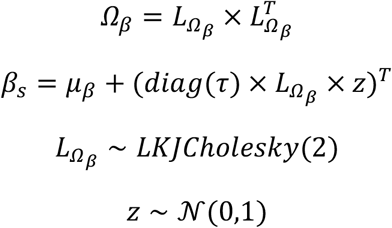

where multiplying the Cholesky factor of the correlation matrix by the standard normally distributed additional parameter *z* and adding the group mean *µ*_*β*_ creates a *β*_*s*_ vector distributed identically to the original model.

While the choice function is identical for each model, the parameters used in generating deck value differ for each. All were fit hierarchically and were modeled with the following priors and hyperpriors:

Rescorla Wagner with a single learning rate (RW1*α*):

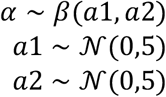

Rescorla Wagner with two learning rates (RW2*α*):

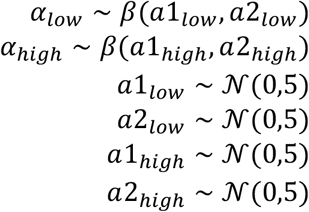

Reduced Bayes with a single hazard rate (RB1*H*):

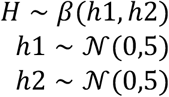

Reduced Bayes with two hazard rates (RB2*H*):

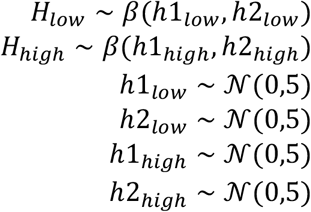

